# Rats spontaneously show categorical responses toward familiar or unfamiliar conspecifics in a habituation-dishabituation task using multiple habituation stimuli

**DOI:** 10.1101/2025.09.25.678521

**Authors:** Shiomi Hakataya, Makiko Kamijo, Genta Toya, Hiroki Koda, Kazuo Okanoya

**Affiliations:** Graduate School of Arts and Sciences, The University of Tokyo, 3-8-1 Komaba, Meguro-ku, Tokyo, 153-8902, Japan; Advanced Comprehensive Research Organization, Teikyo University, 2-21-1 Kaga, Itabashi-ku, Tokyo, 173-0003, Japan; Japan Society for the Promotion of Science, Kojimachi Business Center, 5-3-1 Kojimachi, Chiyoda-ku, Tokyo, 102-0083, Japan; Faculty of Science, University of the Ryukyus, 1 Sembaru, Nishihara-cho, Nakagami-gun, Okinawa, 903-0213, Japan; Faculty of Liberal Arts, Teikyo University, 359 Otsuka, Hachioji-shi, Tokyo, 192-0395, Japan; Research Center for Advanced Science and Technology, The University of Tokyo, 4-6-1 Komaba, Meguro-ku, Tokyo, Japan; School of Environment and Society, Institute of Science Tokyo, 2-12-1 Ookayama, Meguro-ku, Tokyo, 152-8552, Japan; Center for Evolutionary Cognitive Science, The University of Tokyo, 3-8-1 Komaba, Meguro-ku, Tokyo, 153-8902, Japan

**Keywords:** Social familiarity, Rat, Habituation-dishabituation paradigm, Social recognition, Cage-mate

## Abstract

Discriminating between familiar and unfamiliar conspecifics is a fundamental aspect of social recognition. Previous studies have shown that familiarity strongly influences rat social behavior, and that rats can be trained to discriminate between olfactory cues from cage-mates and non-cage-mates in a digging task. However, such discrimination did not occur in a lever-pressing task using live conspecifics as stimuli. In the present study, we investigated whether rats could discriminate social familiarity in a simple, less-demanding and naturalistic context using whole-animal stimuli. We employed a habituation-dishabituation paradigm in which multiple cage-mate or non-cage-mate individuals were presented during the habituation phase. We examined whether rats would habituate to the familiarity category among these multiple stimuli and subsequently show dishabituation when the stimulus class switched (cage-mate to non-cage-mate or vice versa). During the habituation phase, exploratory responses declined over successive sessions, suggesting habituation to the common element among multiple cage-mates or non-cage-mates, i.e., familiarity/unfamiliarity. When all subjects were considered together, clear dishabituation was not observed, indicating individual variation in habituation. However, a post-hoc analysis restricted to subjects showing sufficient habituation revealed significant dishabituation to the stimulus class switch. These findings suggest that rats can spontaneously classify multiple conspecifics according to their familiarity without training. Rats may flexibly employ different forms of social recognition and information such as individual identity or familiarity-based heuristics to guide adaptive social behavior in varying contexts.

**Highlights:**

- We tested familiarity discrimination in rats by habituation-dishabituation task
- Multiple cage-mate or non-cage-mate stimuli were presented during habituation
- Exploratory responses decreased as habituation sessions progressed
- Dishabituation occurred in individuals who exhibited sufficient habituation
- Rats showed categorical responses toward familiarity or unfamiliarity

## 1. Introduction

In social recognition, animals recognize sets of individuals (class-level cognition) or each individual separately (individually recognition) (Wiley et al., 2013; Yorzinski, 2017). The ability to discriminate between familiar and unfamiliar conspecifics known as social familiarity discrimination is an essential form of class-level recognition. Because it is cognitively less demanding than remembering and recognizing each conspecific individually, social familiarity discrimination can be viewed as an adaptive heuristic for managing the complexity of the social environment (Coulon et al., 2011; Jones et al., 2014; Suwandschieff et al., 2023; Wilkinson et al., 2010). Experimental evidence shows some animals can be trained to classify images of conspecifics according to their familiarity (cattle: Coulon et al., 2011; pigeon: Wilkinson et al., 2010).

Laboratory rats can be a good model for studying social familiarity discrimination. Familiarity with a partner can modulate various social behaviors in rats (e.g., Alberts & Galef, 1973; Hackenberg et al., 2021; Kiyokawa et al., 2014; Rogers-Carter et al., 2018; Smith et al., 2015). Jones et al. (2014) demonstrated that rats can be trained to discriminate between olfactory cues from familiar cage-mates and unfamiliar non-cage-mates in a digging task, which uses natural digging behavior into media containing odors as response. However, rats failed to reach the learning criterion in a lever-pressing task using live conspecific stimuli. While learning tasks such as digging can clearly demonstrate stimulus discrimination, they are often time-consuming. Also, the collection and preparation of suitable odor samples require careful handling. Development of a simpler task could broaden opportunities for future research (e.g., Wada et al., 2025), including investigations into mechanisms underlying familiarity discrimination. We therefore aimed to test whether social familiarity discrimination could be achieved in a simpler and semi-natural task without specific training using whole-animal stimuli.

The habituation-dishabituation paradigm is widely used to assess olfactory discrimination and social recognition in rodents (Slotnick et al., 2004). In this task, repeated exposure to stimuli from the same class typically induces a gradual decline in behavioral responses during habituation. A subsequent recovery of responses to stimuli from a different class during dishabituation is generally interpreted as evidence of discrimination. However, if only a single stimulus is repeatedly presented during habituation, it remains possible that the recovered responses during dishabituation reflect discrimination between two individual stimuli rather than between two stimulus classes. Presenting multiple stimuli from the same class during habituation allows assessment of discrimination based on shared features across stimuli (Kondo et al., 2005). Accordingly, in our paradigm, we presented multiple cage-mates or non-cage-mates during the habituation phase to examine whether rats would show habituation and dishabituation to the common factor of familiarity versus unfamiliarity.

In this study, we presented cage-mate and non-cage-mate stimuli and measured the duration of subjects’ social exploration. In the cage-mate habituation condition, animals were presented with multiple cage-mate stimuli during habituation and non-cage-mates during dishabituation; vice versa for the non-cage-mate habituation condition. We predicted that rats would habituate to the common factor among cage-mates or non-cage-mates (i.e., familiarity or unfamiliarity), and would show dishabituation when the stimulus class switched to the other one.

## 2. Materials and Methods

More details are provided in Supplementary Methods.

### 2.1. Ethical statements

All procedures were conducted in accordance with the experimental implementation regulations of the University of Tokyo and were approved by the Animal Experimental Committee at the University of Tokyo, Graduate School of Arts and Sciences (Permission Number: 2022-5).

### 2.2. Animals

Subjects were 32 male Sprague-Dawley rats (SLC Japan Inc., Hamamatsu, Japan). Individuals housed in the same cage (four per cage) were designated as cage-mates. Experiments were conducted 5 weeks after the start of co-housing, when the rats were 17 weeks old. Prior to this study, all rats experienced handling, a series of behavioral tests and group tracking in a large cage. Because none of these prior procedures resembled the experimental settings or protocols used here, they were unlikely to have affected the results.

Subjects were kept in a room under a 12:12 h light/dark cycle (lights on at 07:00). All experiments were conducted during the light period. Each group received 125 g of standard laboratory chew daily (Lab Diet, PMI Nutrition International, St. Louis, USA). Water was freely available.

### 2.3. Apparatus

Experiments were conducted in a custom-made open field (80 × 80 × 50 cm). Stimulus animals were presented in black acrylic cages (18 cm length × 25.5 cm width × 52 cm height). The lower half of the front side of each cage was covered with 5 × 5 mm acrylic rods, allowing interactions between the subject and the stimulus animal. The stimulus cage was placed centrally along the back wall of the arena (**Figure 1a**). A camera (HERO9 BLACK, GoPro) was mounted above the arena for video recording.

**Figure 1.**
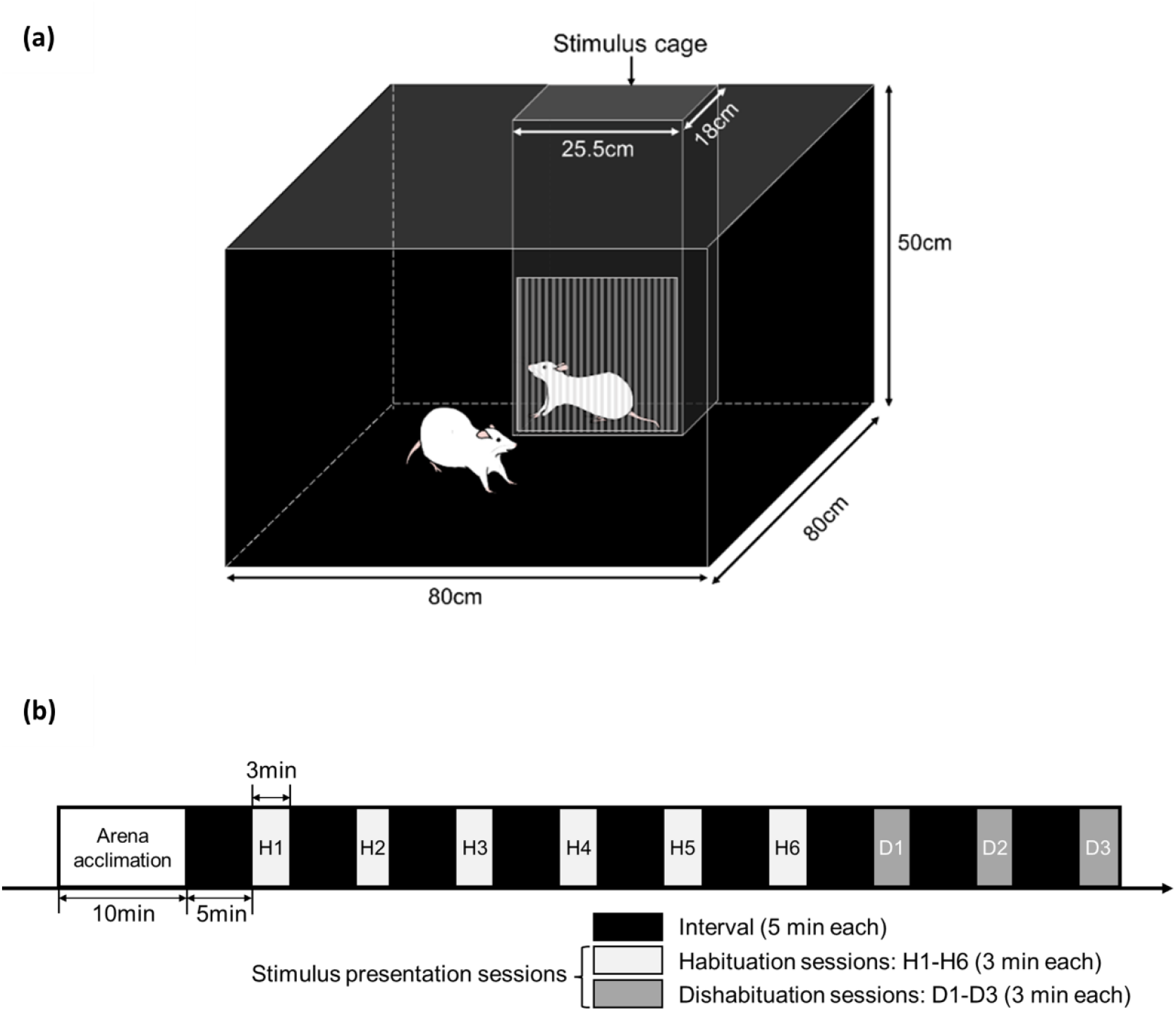
Experimental procedures. (a) The experimental apparatus. The apparatus was a black acrylic open field. Stimulus individuals were presented in black acrylic cages placed centrally along the back wall of the arena. The lower half of the front side of each cage was covered with acrylic rods and allowed social interactions between the subject and the stimulus animal. (b) Habituation-dishabituation paradigm. During the arena acclimation phase, subjects were introduced to the arena and allowed to explore freely for 10 minutes. Each stimulus presentation session lasted for 3 minutes. The inter-session interval was 5 min. During habituation sessions (H1-H6), three cage-mate or non-cage-mate stimuli were each presented twice in a pseudo-random order. In dishabituation sessions (D1-D3), three different cage-mate or non-cage-mate stimuli were each presented once in a random order.

### 2.4. Experimental procedures

The habituation-dishabituation task included the two conditions: the cage-mate habituation (CMH) and the non-cage-mate habituation (NCMH) condition. Each subject was randomly assigned to one of these conditions. In the CMH condition, animals were presented with cage-mate stimuli during habituation and non-cage-mates during dishabituation; vice versa in the NCMH condition. In both conditions, the cage-mate stimuli were three individuals from the subject’s own group, and the non-cage-mate stimuli were three individuals from a different group. All individuals served as both subjects and stimulus animals.

The behavioral task consisted of an arena acclimation phase followed by stimulus presentation (6 habituation sessions and 3 dishabituation sessions) (**Figure 1b**). During arena acclimation, subjects were introduced to the arena and allowed to explore freely for 10 minutes. In each stimulus presentation session, a cage-mate or non-cage-mate stimulus was presented for 3 minutes. The inter-session interval was 5 minutes, during which the apparatus was cleaned using 75% ethanol.

In habituation sessions, the three cage-mate or non-cage-mate stimuli were each presented twice in a pseudo-random order, ensuring that the same stimulus was not presented in two consecutive sessions. In dishabituation sessions, the three cage-mate or non-cage-mate stimuli were each presented once in a random order. The testing order of subjects was randomized.

### 2.5. Analysis

Exploratory responses to stimuli were quantified as the total time the subject’s nose remained on the front of the stimulus cage or within 1.5 cm of it (nose length: approximately 1.5 cm). This was measured using DeepLabCut 2.3 (Mathis et al., 2018; Nath et al., 2019). Due to mechanical troubles, all sessions for two individuals and four habituation sessions from four different individuals were excluded from analysis.

We analyzed the data by fitting generalized linear mixed models (GLMMs) in R 4.5.0 (R Core Team, 2025; glmmTMB: Brooks et al., 2017). All models included the duration of responses as the response variable, with session, condition and their interaction as fixed effects, and subject ID as a random effect.

## 3. Results

We analyzed whether exploratory responses changed during habituation sessions using GLMM (**Figure 2a**). The parameter coefficient of session was estimated as a negative value (*β* ± *SE* = -0.161 ± 0.0308, *z* = -5.22, *p* < 0.001, 95% CI [-0.221, -0.0996]). The fixed effect of condition (*β* ± *SE* = -0.0845 ± 0.196, *z* = -0.430, *p* = 0.667) and the session × condition interaction (*β* ± *SE* = 0.0190 ± 0.0426, *z* = 0.445, *p* = 0.656) were not significant. These results indicate that exploratory responses decreased as habituation sessions progressed and that subjects habituated to stimuli within the same class (cage-mate or non-cage-mate).

**Figure 2.**
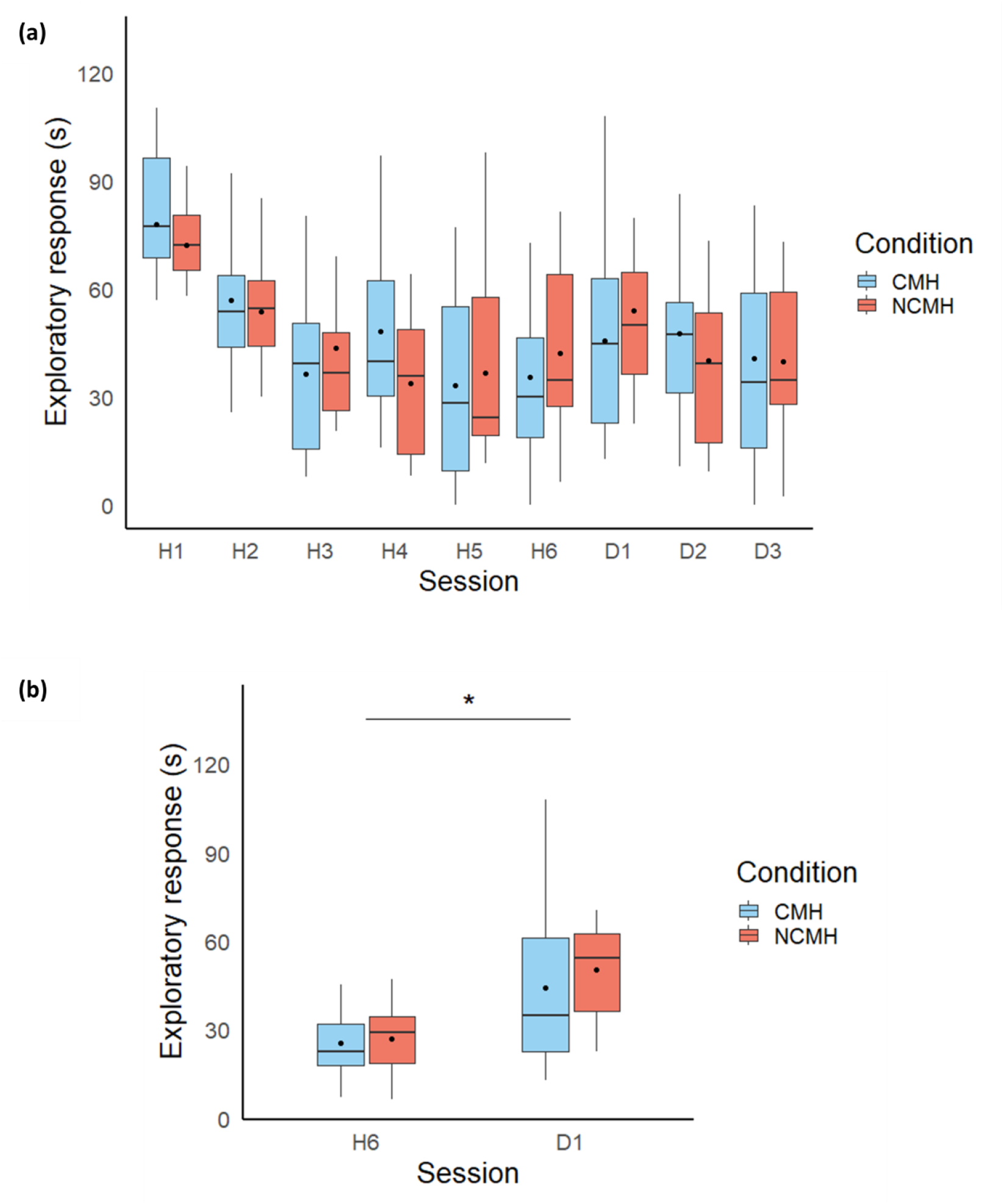
Exploratory responses toward cage-mates and non-cage-mates. Box plots represent exploratory responses toward cage-mate and non-cage-mate stimuli. Blue indicates the cage-mate habituation condition (CMH) and red, the non-cage-mate habituation condition (NCMH), respectively. Black circles represent mean values. (a) whole data (N = 30; N = 15 from the CMH condition, N = 15 from the NCMH condition). (b) only individuals who showed sufficient habituation (N =21/30; N = 11/15 from the CM condition, N = 10/15 from the NCM condition). * indicates *p* < 0.05.

We next compared exploratory responses between the last habituation session and first dishabituation session to examine whether dishabituation occurred (**Figure 2a**). Neither the fixed effect of session (*β* ± *SE* = 0.182 ± 0.215, *z* = 0.847, *p* =0.397) nor condition (*β* ± *SE* = 0.106 ± 0.219, *z* = 0.484, *p* = 0.629) nor the session × condition interaction (*β* ± *SE* = 0.0678 ± 0.299, *z* = 0.227, *p* = 0.821) was significant. These results suggest subjects did not show dishabituation in response to the stimulus class switch.

Further inspection of the data revealed substantial individual variability in responses during habituation sessions, suggesting that habituation had not been fully induced in some individuals. Therefore, as an additional post-hoc analysis, we examined whether dishabituation occurred in those that had habituated (N =21/30 in total; N = 11/15 from the CM condition, N = 10/15 from the NCM condition) (**Figure 2b**). We operationally defined individuals as sufficiently habituated if they met both of the following criteria: (1) the response in the last habituation session was less than two-thirds of that in the first habituation session, and (2) the average response during the fourth to sixth habituation sessions was lower than that during the first to third habituation sessions.

When responses were compared between the last habituation session and the first dishabituation session in this subset, the fixed effect of session was significant (*β* ± *SE* = 0.554 ± 0.226, *z* = 2.45, *p* = 0.0145, 95% CI [0.0967, 1.01]), whereas the fixed effect of condition (*β* ± *SE* = 0.0618 ± 0.232, *z* = 0.267, *p* = 0.790) and the session × condition interaction (*β* ± *SE* = 0.0649 ± 0.324, *z* = 0.200, *p* = 0.841) were not significant. These results suggest that exploratory responses increased from the last habituation session to the first dishabituation session, indicating that dishabituation occurred in individuals that had been adequately habituated during the habituation phase.

## 4. Discussion

In this study, we examined social familiarity discrimination in rats using a habituation-dishabituation paradigm. We presented multiple cage-mate or non-cage-mate stimuli during the habituation phase and then tested whether rats would show dishabituation when the stimulus class switched to the other one. During the habituation phase, exploratory responses decreased as sessions progressed. Clear dishabituation to the stimulus class switch was not observed when all individuals were pooled. However, the post-hoc analysis restricted to subjects that habituated sufficiently revealed significant dishabituation.

Our results suggest that rats spontaneously exhibited a categorical response toward familiarity or unfamiliarity in the current experimental setting and that rats can classify conspecifics according to their familiarity. Insufficient habituation in some subjects may have resulted from using live conspecific stimuli, whose activity or social relationships with the subjects could have obscured responses. To control for these factors, it may be useful to restrain stimulus animals’ movement and increase the variety of stimulus animals. Also, in the habituation-dishabituation paradigm, individual differences in the degree of habituation can be large (Baciadonna et al., 2019; Charlton et al., 2007). In this study, we kept the number of habituation sessions consistent across subjects to control the amount of stimulus exposure. Future studies could increase exposure to habituation stimuli to ensure sufficient habituation in all individuals.

The habituation-dishabituation paradigm with multiple conspecific stimuli is easy to implement and may be a promising task for investigating social familiarity recognition and its underlying mechanisms (c.f. mouse: Wolf et al., 2024; sheep: Kendrick et al., 2001). The specific cues by which rats differentiate cage-mates from non-cage-mates remain unknown. Jones et al. (2014) proposed that there may be group-specific odors (c.f. chemical cues of familiarity; ground squirrel: Hare, 1994; bat: Rizzi et al., 2025), or that rats may form categories of familiarity and unfamiliarity. Regarding odor cues, extended cohabitation may lead to convergence of odors among group members through social interactions and the sharing of resources such as food and a home base. One possible mechanism includes the transmission of gut microbiota among group members through interactions (Raulo et al., 2021), as bacteria are known to influence the discriminability of urine odors in rats (Schellinck et al., 1991).

It remains to be investigated whether rats utilize different types of social recognition, such as individual recognition (Gheusi et al., 1997; Hopp et al., 1985) and the use of familiarity heuristics, both of which may guide adaptive social behavior depending on the situation. For instance, rats may rely on individual recognition when directly interacting with a small number of individuals (Hakataya et al., 2023; Proops et al., 2021). However, in the wild, where there may be many neighboring individuals (Davis, 1953) and odor cues may be degraded, familiarity discrimination could be a more efficient way for managing frequent encounters.

## Data availability Statement

Data and analysis codes are available from the following OSF project: https://osf.io/apngb/?view_only=5dd27745b1f54c92bc75f42f4e8b0360.

## CRediT authorship contribution statement

- Shiomi Hakataya: Conceptualization, Methodology, Formal analysis, Investigation, Resources, Writing - Original Draft, Visualization, Funding acquisition
- Makiko Kamijo: Conceptualization, Methodology, Writing - Review & Editing, Funding acquisition
- Genta Toya: Methodology, Software, Writing - Review & Editing, Funding acquisition
- Hiroki Koda: Resources, Writing - Review & Editing, Funding acquisition
- Kazuo Okanoya: Writing - Review & Editing, Supervision, Funding acquisition

## Acknowledgements

We thank Reo Wada and Kosuke Yoshida for their assistance in animal care. We are also grateful to the members of Okanoya laboratory at Teikyo University for stimulating discussion.

## Declaration of Generative AI and AI-assisted Technologies in the Writing Process

During the preparation of this work, the first author used ChatGPT in order to improve the readability and language of the manuscript. After using this tool, the authors reviewed and edited the content as needed and take full responsibility for the content of the published article.

