## Supplementary Methods for "Rats spontaneously show categorical responses toward familiar or unfamiliar conspecifics in a habituation-dishabituation task using multiple habituation stimuli"

Data, analysis codes, and instructions are available from the following OSF project. https://osf.io/apngb/?view_only=5dd27745b1f54c92bc75f42f4e8b0360

#### 1. Subjects and housing

Subject were 32 male Sprague-Dawley (SD) rats. Rats were 8 weeks old at the time of purchase and initially housed in pairs. Starting from approximately 12 weeks of age, two previously housed pairs were combined to form cages of four individuals each. Individuals housed in the same cage were designated as cage-mates. Experiments were conducted 5 weeks after the start of co-housing, when the rats were 17 weeks old.

To acclimate subjects to the experimenter, each rat was handled for 5 minutes on five separate days at 9 weeks of age. Subjects also experienced a series of behavioral tests, including: the glove test (tameness), open field test (activity level), novel object test (novelty exploration), elevated plus maze test (anxiety), and food competition test (social dominance), as well as group tracking in a large cage. Because these prior procedures did not resemble the experimental settings or protocols used here, they were unlikely to have affected the results.

Subjects were kept in a room under a 12:12 h light/dark cycle (lights on at 07:00 am). All experiments were conducted during the light period. To pervert overeating, each group received 125 g of standard laboratory chew daily (approximately 31 g per animal). Water was freely available.

#### 2. Apparatus

Experiments were conducted in a black acrylic open field (80 × 80 × 50 cm) illuminated by an LED light above the center (approximately 430 lx at center, 380 lx at corners) to provide even lighting. Stimulus animals were presented in black acrylic cages (18 cm length × 25.5 cm width × 52 cm height). The lower half of the front side of each cage was covered with 5 × 5 mm acrylic rods spaced 5 mm apart, allowing interactions such as sniffing between the subject and the stimulus animal. The stimulus cage was placed centrally along the back wall of the arena. A camera (HERO9 BLACK, GoPro) was mounted above the arena for video recording.

#### 3. Experimental procedure

##### 3.1. Task design

The habituation-dishabituation task included the two conditions: the cage-mate habituation (CMH) and the non-cage-mate habituation (NCMH) condition. Each subject was randomly assigned to one of these conditions. In the CMH condition, animals were presented with cage-mate stimuli during the habituation phase and non-cage-mates during the dishabituation phase; vice versa in the NCMH condition. In both conditions, the cage-mate stimuli were three individuals from the subject's own group, and the non-cage-mate stimuli were three individuals from a different group. All individuals served as both subjects and stimulus animals. Stimulus assignments were quasi-random, ensuring each rat experienced the stimulus role equally.

#### 3.2. Task flow

Prior to testing, animals were acclimated to the stimulus cage twice. Each habituation session lasted 5 minutes, during which 10 rice-puffs were placed on the cage floor to promote acclimation. In the second session, the stimulus cage was placed in the arena, and we confirmed that all rice-puffs had been eaten.

For each subject's experiment, two groups were used: the group to which the subject belonged and the group to which the stimulus animals belonged. Immediately prior to the experiment, eight individuals belonging to these two groups were transferred to individual small cages, transported to the experimental room, and isolated for 15 minutes. These procedures controlled for acute responses to separation from cage-mates and exposure to a novel environment.

The behavioral task consisted of an arena acclimation phase followed by stimulus presentation (6 habituation sessions and 3 dishabituation sessions). During the arena acclimation, subjects were introduced to the center of the front side of the arena and allowed to explore freely for 10 minutes. In each stimulus presentation session, a cage-mate or non-cage-mate stimulus was presented for 3 minutes. The inter-session interval was 5 minutes, during which the apparatus was cleaned using 75% ethanol.

In habituation sessions, the three cage-mate or non-cage-mate stimuli were each presented twice in a pseudo-random order, ensuring that the same stimulus was not presented in two consecutive sessions. In dishabituation sessions, the three cage-mate or non-cage-mate stimuli were each presented once in a random order. The testing order of subjects was randomized. When multiple subjects were tested on the same day, one-hour interval was maintained between experiments to allow social interactions in the home cage.

### 4. Analysis

#### 4.1. Video analysis

Exploratory responses to stimuli were quantified as the total time the subject's nose remained on the front of the stimulus cage or within 1.5 cm of it. The 1.5-cm threshold was based on the apparent size of a rat's nose in videos (approximately 1.5cm), since rats typically keep their noses very close to stimuli during exploratory sniffing.

The videos were analyzed using DeepLabCut 2.3 (CPU: Intel® Xeon® W-2123; GPU: NVIDIA GeForce RTX 2080 Ti; OS: Ubuntu 20.04 LTS), and XY coordinate data for body parts were obtained. The coordinate data was further analyzed using a Python program, which automatically calculated the total time spent in, and the number of entries into, the designated area. Estimated coordinate data with a likelihood > 0.8 were used for the calculations. Due to mechanical troubles, all sessions for two individuals and four habituation sessions from four different individuals were excluded from analysis.

### 4.2. Statistical analysis

We analyzed the data by fitting generalized linear mixed models (GLMMs) in R 4.5.0.

#### 4.2.1. Habituation

To analyze whether exploratory responses changed across habituation sessions, a GLMM was constructed using glmmTMB function of glmmTMB package:

*glmmTMB (duration ~ session * condition + (1|ID), family = Gamma (link = "log"))*

- duration: the total time of exploratory responses to the stimulus
- condition: CMH (0) or NCMH (1)
- session: habituation sessions (1 - 6)
- (1|ID): random intercept for each subject

Although the data roughly followed the gamma distribution, zeros occurred in 3 out of 176 observations. Because these accounted for a small fraction of the data, these were omitted.

#### 4.2.2. Dishabituation

To examine whether dishabituation occurred, a GLMM was constructed:

*glmmTMB (duration ~ session * condition + (1|ID), family = Gamma (link = "log"))*

- duration: the total time of exploratory responses to the stimulus
- condition: CMH (0) or NCMH (1)
- session: the last habituation session (6) and first dishabituation session (7) as factor levels
- (1|ID): random intercept for each subject

To analyze whether dishabituation occurred in subjects that showed sufficient habituation, the same analysis was applied to this subset. Omitted zeros accounted for a small fraction of the data (1/60 observation in the whole dishabituation data; 1/42 observations in the subset data).
